# Comprehensive analysis of fungal G1 cyclin docking motif sequences that control CDK regulatory potency in vivo

**DOI:** 10.1101/2020.03.02.973354

**Authors:** Sushobhana Bandyopadhyay, Samyabrata Bhaduri, Mihkel Örd, Norman E. Davey, Mart Loog, Peter M. Pryciak

## Abstract

Cyclin-dependent kinases (CDKs) control the ordered series of events during eukaryotic cell division. The stage at which individual CDK substrates are phosphorylated can be dictated by cyclin-specific docking motifs. In budding yeast, substrates with Leu/Pro-rich (LP) docking motifs are recognized by Cln1/2 cyclins in late G1 phase, yet the key sequence features of these motifs and the conservation of this mechanism were unknown. Here we comprehensively analyzed LP motif requirements in vivo by combining a competitive growth assay with mutational scanning and deep sequencing. We quantified the impact of all single-residue replacements in five different LP motifs, using six distinct G1 cyclins from diverse fungi including medical and agricultural pathogens. The results reveal the basis for variations in potency among wild-type motifs, and allow derivation of a quantitative matrix that predicts the potency of other candidate motifs. In one protein, Whi5, we found overlapping LP and phosphorylation motifs with partly redundant effects. In another protein, the CDK inhibitor Sic1, we found that its LP motif is inherently weak due to unfavorable residues at key positions, and this imposes a beneficial delay in its phosphorylation and degradation. The overall results provide a general method for surveying viable docking motif sequences and quantifying their potency in vivo, and they reveal how variations in LP motif potency can tune the strength and timing of CDK regulation.

## INTRODUCTION

Protein phosphorylation regulates a wide variety of cellular processes, and eukaryotic cells possess hundreds of different protein kinases. For these numerous kinases to perform distinct regulatory functions, they cannot all phosphorylate the same substrates. Instead, several types of mechanism allow protein kinases to be selective [1, 2]. These include the recognition of preferred phospho-acceptor residues and their surrounding sequence context by the catalytic site in the enzyme, as well as mechanisms that increase the physical association and/or co-localization of the kinase with its substrate. Among the latter type of mechanism, several kinases (or their partner proteins) recognize “docking motifs” in the substrate, which are short linear motifs (SLiMs) that are separate from the phosphorylated residues, often in disordered regions [1-3]. Because such motifs are recognized by only a subset of kinases, they enhance the specificity of phosphorylation. Similar mechanisms regulate the ability of phosphatases to catalyze the reverse reactions, which helps ensure that protein phosphorylation is dynamic [4-6]. There is a pressing need for better understanding of how docking interactions control phosphorylation networks, and the rules that govern whether a particular substrate will be efficiently engaged by a given kinase. Yet, for many docking motifs there is a lack of comprehensive information about the range of possible sequences that can be recognized and distinguished from the thousands of other competing peptide sequences in vivo.

Cyclin-dependent kinases (CDKs) are the central regulators of eukaryotic cell division [7]. All eukaryotes have evolved multiple cyclin-CDK complexes that function at different stages of the cell cycle, and therefore it is important to elucidate the molecular basis for their specializations. In the budding yeast *Saccharomyces cerevisiae*, a single CDK enzyme (Cdk1) associates with multiple different cyclins, which impart distinct properties to the enzyme complex as the cell cycle proceeds [8, 9]. Recent studies indicate that many of these cyclins recognize unique docking motifs in substrates that allow them to be phosphorylated in a cyclin-specific manner [10-16]. We have been studying docking interactions involving G1 cyclins in yeast. This process involves a SLiM called an “LP” motif (for its enrichment in Leu and Pro residues), which is recognized specifically by the late-G1 cyclins (Cln1 and Cln2), and not by the early-G1 cyclin (Cln3) or by the later S- and M-phase cyclins (Clb1-Clb6). Previous studies indicate that these LP docking interactions increase the efficiency, specificity, and multiplicity of substrate phosphorylation [10, 11, 17]. Nevertheless, several fundamental issues remain poorly understood, including the key features of the LP motif sequence, their prevalence in the proteome, and the degree of conservation of LP docking among distinct organisms and cyclin subtypes.

Prior evidence suggests that functional LP motifs exist in multiple Cln2-CDK substrates, as their phosphorylation in vitro can be inhibited by competing LP motif peptides or by Cln2 mutations that disrupt LP motif recognition [11, 17]. Yet in only three proteins has a specific LP motif been identified as responsible: Sic1 (a CDK inhibitor), Ste5 (a MAPK scaffold protein), and Ste20 (a membrane kinase) [10, 11, 17]. While these identified motifs all share a minimal LxxP sequence, the key recognition features of LP motifs remain poorly defined, which hinders their identification in known or candidate substrates and obscures insight into the basis of cyclin specificity. Moreover, in addition to Cln3 and Cln1/2 cyclin types, some budding yeast species contain a third type of G1 cyclin, called Ccn1, which is also capable of recognizing LP motifs [17]. In contrast, most non-yeast fungi have only a single G1 cyclin (hereafter, “Cln”), which likely diverged prior to the expansion of the budding yeast genes into the three G1 types [18]. Thus, it is unclear if LP docking is an ancient function that is conserved broadly among fungal G1 cyclins or if it evolved more recently only in yeasts.

In this study we investigated the conservation of the LP docking mechanism and the sequence features of LP motifs that dictate their functional potency in vivo. We report that LP motifs can be recognized by a broad variety of G1 cyclins throughout the fungal kingdom, rather than being limited to budding yeasts. We performed a comprehensive analysis of LP motifs, involving systematic mutational scanning and an in vivo competitive growth assay, to derive a quantitative measure of the functional impact of all substitutions at all positions in the linear sequence. These efforts revealed the key sequence features that control the recognition of LP motifs, and how sequence variations can modulate the strength and timing of CDK regulation.

## RESULTS

### LP motif recognition is broadly conserved among fungal G1 cyclins

Previously, we found that LP motifs are recognized by two types of G1 cyclins from budding yeast, Cln1/2 and Ccn1, but not by a third type, Cln3 [17]. Most other fungi have only one type of G1 cyclin (Figure 1A), for which there was minimal functional information and none about docking. Thus, we used our prior in vivo phosphorylation assay to test if G1 cyclins from these more distant fungi can recognize LP motifs. The foreign cyclin genes were inserted into a *P*_*GAL1*_ vector that permits galactose-inducible expression in *S. cerevisiae*, and also adds an N-terminal GST tag plus one half of a leucine zipper sequence; this latter feature allows the cyclin to engage a control substrate (with the partner half zipper), so that its function can be measured independent of LP motif recognition [17, 19]. We chose representative cyclins from the three major subdivisions of Ascomycota and from three genera of Basidiomycota (Figure 1A-B). Each was tested for the ability to drive phosphorylation of a protein fragment with no docking sequence, with an LP motif, or with a half leucine zipper (Figure 1B). As seen previously [17, 19], of two *S. cerevisiae* cyclins, only Cln2 and not Cln3 promoted phosphorylation of the substrate with the LP motif. Remarkably, most other cyclins behaved like Cln2, in that they could use the LP motif to promote substrate phosphorylation. The sole exception was Puc1 from the fission yeast *S. pombe*, but it also showed weak expression and functioned poorly with the leucine zipper substrate. In contrast, a Puc1 cyclin from another fission yeast, *S. japonicus*, was able to drive phosphorylation of the LP motif substrate, and hence the *S. pombe* cyclin was an outlier.

**FIGURE 1:**
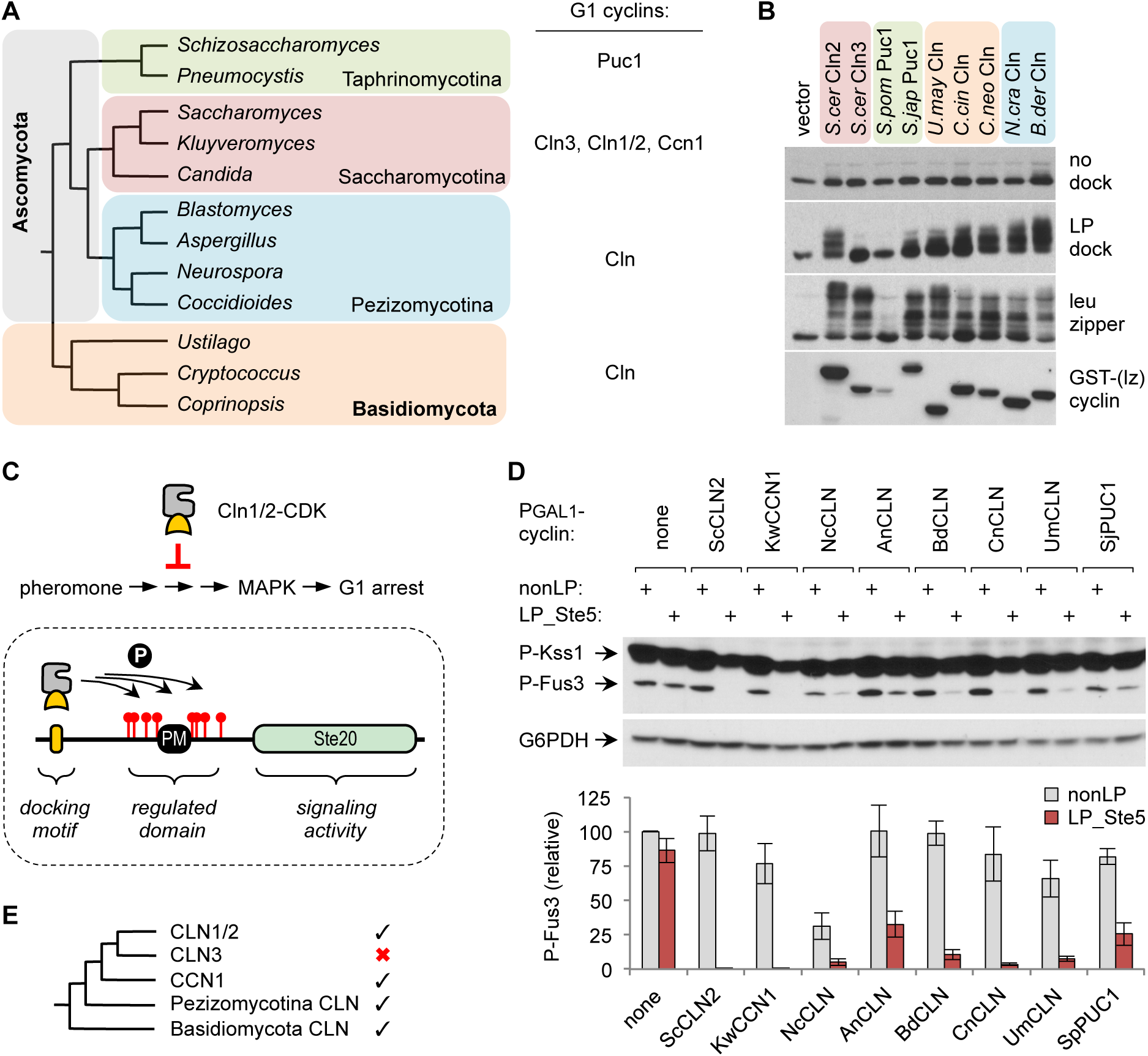
Conservation of LP docking ability among fungal G1 cyclins. (A) Phylogenetic tree of representative fungal genera in Ascomycota and Basidiomycota, and the types of G1 cyclins found in the indicated subdivisions. (B) Fungal G1 cyclins were expressed in *S. cerevisiae* as galactose-inducible fusions to GST plus a half leucine zipper (GST-[lz]). The cells also harbored HA-tagged substrates with no docking site, or an LP docking motif, or the partner half leucine zipper. Reduced mobility indicates phosphorylation. (C) *Top*, Cln1/2-CDK activity inhibits pheromone-induced MAPK activation and G1 arrest. *Bottom*, the Ste20^Ste5PM^ chimera contains a plasma membrane-binding domain (PM) and flanking CDK sites from Ste5, which can be phosphorylated and inhibited if a functional LP docking motif is present. (D) *P*_*GAL1*_*-cyclin* strains harboring Ste20^Ste5PM^ chimeras with either a functional LP motif (LP_Ste5) or a control sequence (nonLP) were treated with pheromone (50 nM, 15 min.) and tested for MAPK phosphorylation. Below are quantified results (mean ± SEM, n = 4). (E) Phylogeny of fungal G1 cyclins [18] compared with observed docking abilities suggests that LP motif recognition is an ancient, conserved function that was lost by the Cln3 group.

The conservation of LP motif recognition was confirmed via a separate assay that is based on the ability of Cln2 to inhibit signaling in the yeast pheromone response pathway [20, 21] (Figure 1C, top). The ordinary target of this inhibition is the scaffold protein Ste5 [21], but the regulatory effects can be transferred to a chimeric signaling protein (Ste20^Ste5PM^; Figure 1C, bottom; [10]), which provides a useful setting for testing candidate LP motifs. Using this chimera, we compared *P*_*GAL1*_*-cyclin* constructs for their ability to inhibit pheromone signaling (Figure 1D). In addition to *CLN2* and *K. waltii CCN1*, all of the more distantly related G1 cyclins could inhibit activation of Fus3, the pheromone pathway MAPK. The magnitude of inhibition was weaker for some cyclins (e.g., from *A. nidulans* or *S. japonicus*), but all showed a clear dependence on a functional LP motif. (The *N. crassa* cyclin was partly inhibitory even without an LP motif, implying a non-specific or toxic effect, which we did not study further as it did not impact other experiments.) Collectively, these results indicate that recognition of LP motifs is an ancient property of fungal G1 cyclins that remains broadly intact among extant forms throughout the subkingdom Dikarya. Furthermore, the conservation pattern suggests that this function was lost by Cln3 cyclins after they diverged from the Cln1/2 group (Figure 1E).

### Extended LP motif sequences impart variation in cyclin recognition

The sequence features that define a functional LP motif are poorly defined. Of the three motifs known to function in the context of the native protein, those from Sic1 and Ste5 share a 4-residue sequence, LLPP (Figure 2A); however, the key residues in this stretch and the role of flanking sequence were unknown, and they are only partly similar to the sequence in Ste20 (LDDP). To begin defining the required features, we mutated the Ste5 motif in blocks of four residues around the LLPP sequence (Figure 2B). Residues on the N-terminal side were dispensable, whereas the LLPP sequence itself and the next four C-terminal residues were needed for recognition by both Cln1/2 and Ccn1, which agrees with our previous finding of a role for residues C-terminal to the LDDP motif in Ste20 [10]. Further evidence implicated these C-terminal residues in differential recognition by distinct cyclins. Namely, despite a shared LLPP sequence, the Ste5 motif was recognized by both Cln1/2 and Ccn1 whereas the Sic1 motif was recognized only by Cln1/2 (Figure 2C). These behaviors could imply that Cln1/2 and Ccn1 have distinct preferences in the C-terminal part of the motif, or that LP recognition by Ccn1 is inherently weaker and hence is sensitized to weak motifs. In either case, they suggest a basis by which substrates might be engaged to different extents by different cyclins. Indeed, in binding assays, Ccn1 co-precipitated with only a subset of Cln2 partners, and this subset was distinct from those that bound Cln3 (Figure 2D). Overall, these results suggest that binding sites for both Cln1/2 and Ccn1 extend beyond the core LxxP motif, and that the two groups are differentially sensitive to the residue identities in this flanking region.

**FIGURE 2:**
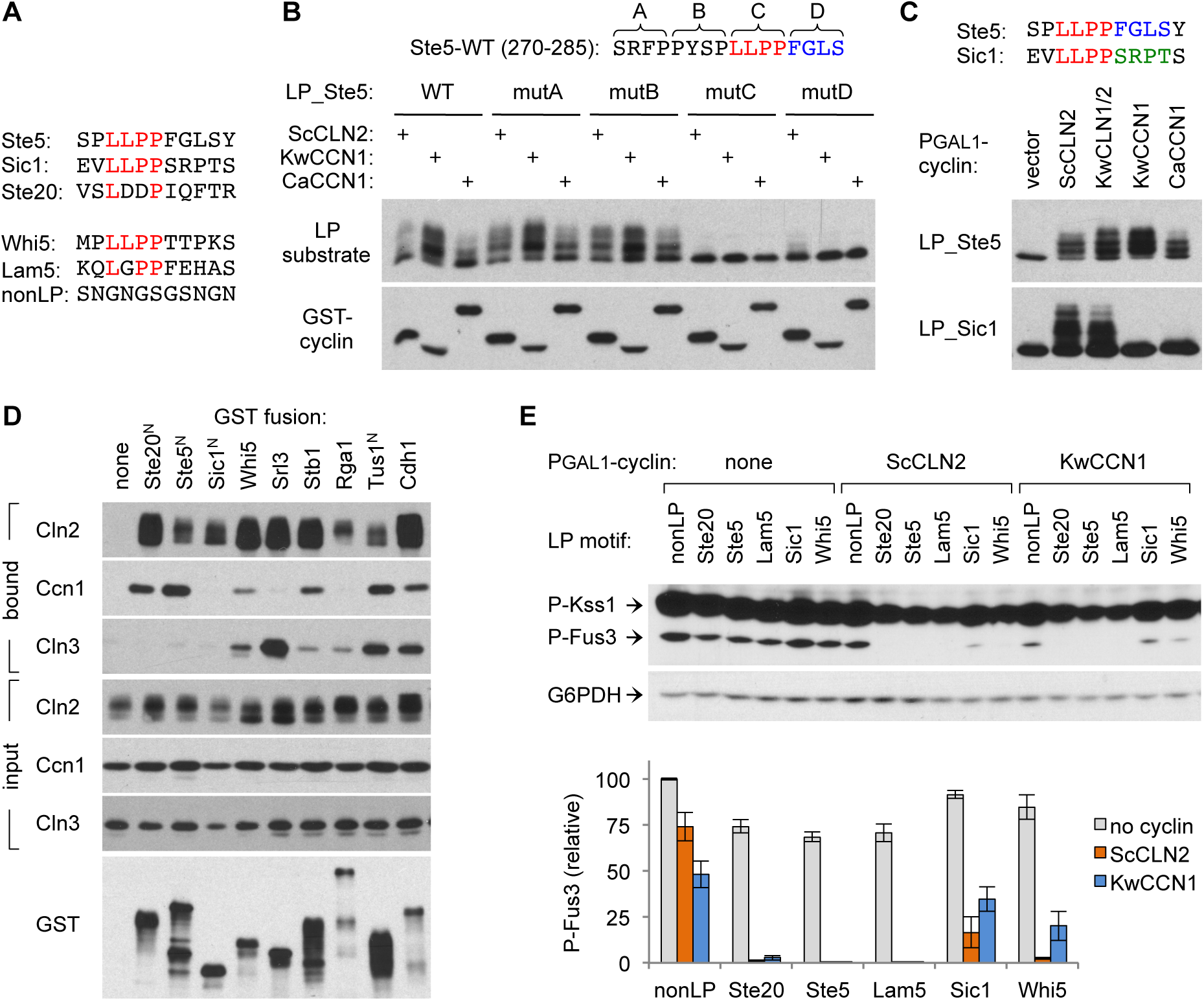
Extended LP motif sequences impart variation in cyclin recognition. (A) LP motif sequences used throughout this study. (B) Four blocks of 4 residues in the Ste5 LP motif region were mutated (to AAAA) and tested for effects on substrate phosphorylation (as in Figure 1B) induced by Cln2 or Ccn1 cyclins. (C) Cln1/2 and Ccn1 type cyclins were tested for phosphorylation of substrates with LP motifs from either Ste5 or Sic1. At top is a comparison of the two LP motif sequences. (D) *P*_*GAL1*_*-cyclin* strains harboring Ste20^Ste5PM^ chimeras with the indicated LP motifs were treated with pheromone (50 nM, 15 min.) and tested for MAPK phosphorylation. Below are quantified results (mean ± SEM, n = 4). (E) G1 cyclins show distinct patterns of binding preference. GST fusions to various Cln2 binding proteins (full-length or N-terminal fragments) were tested for co-precipitation of V5-tagged forms of ScCln2, KwCcn1, or ScCln3. Bound and input fractions are shown, with a representative example of the bound GST signal.

We used the pheromone response inhibition assay to compare a larger set of LP motifs, by making Ste20^Ste5PM^ chimeras with six different docking sequences. These included a nonfunctional control sequence (nonLP), plus LP motifs from Ste5, Sic1, and Ste20, and an LLPP-containing motif from another Cln1/2-regulated substrate, Whi5 (Figure 2A). A final LP sequence was taken from Lam5, which we identified in a recent screen for proteins that bind wild-type Cln2 but not a docking-defective mutant (unpublished observations); the responsible binding motif in Lam5 was of interest here because it contains further sequence variations in both the LxxP and adjacent regions (Figure 2A). We found that all five LP motifs allowed Cln2 to inhibit signaling, although inhibition was noticeably weaker with the Sic1 motif (Figure 2E). With Ccn1, three motifs allowed strong inhibition (Ste5, Ste20, and Lam5), whereas the Whi5 motif was less potent and the Sic1 motif was nearly indistinguishable from the control. Thus, differences in both motif and cyclin can contribute to variation in the strength of CDK regulation.

### A competitive growth assay for the functional potency of cyclins and LP motifs

We wished to systematically interrogate the sequence features of LP motifs that control their recognition and potency. To facilitate this, we designed an in vivo competitive growth assay in which LP motifs provide a selective advantage (Figure 3A). Namely, because pheromone signaling ordinarily causes arrest of cell division, the ability of a cyclin-CDK complex to inhibit signaling can provide a growth advantage. Moreover, because this inhibition depends on the cyclin-LP interaction, strong LP motifs should be more advantageous than weak motifs. Thus, using the Ste20^Ste5PM^ chimera, we compared the relative enrichment or depletion of the five wild-type LP motifs described earlier (Figure 3B). A mixed population of cells harboring these constructs was treated first with galactose (to induce *P*_*GAL1*_*-cyclin* expression) and then with pheromone for various times. The population frequency of each motif was measured via deep sequencing of plasmid DNAs recovered at each time point.

**FIGURE 3:**
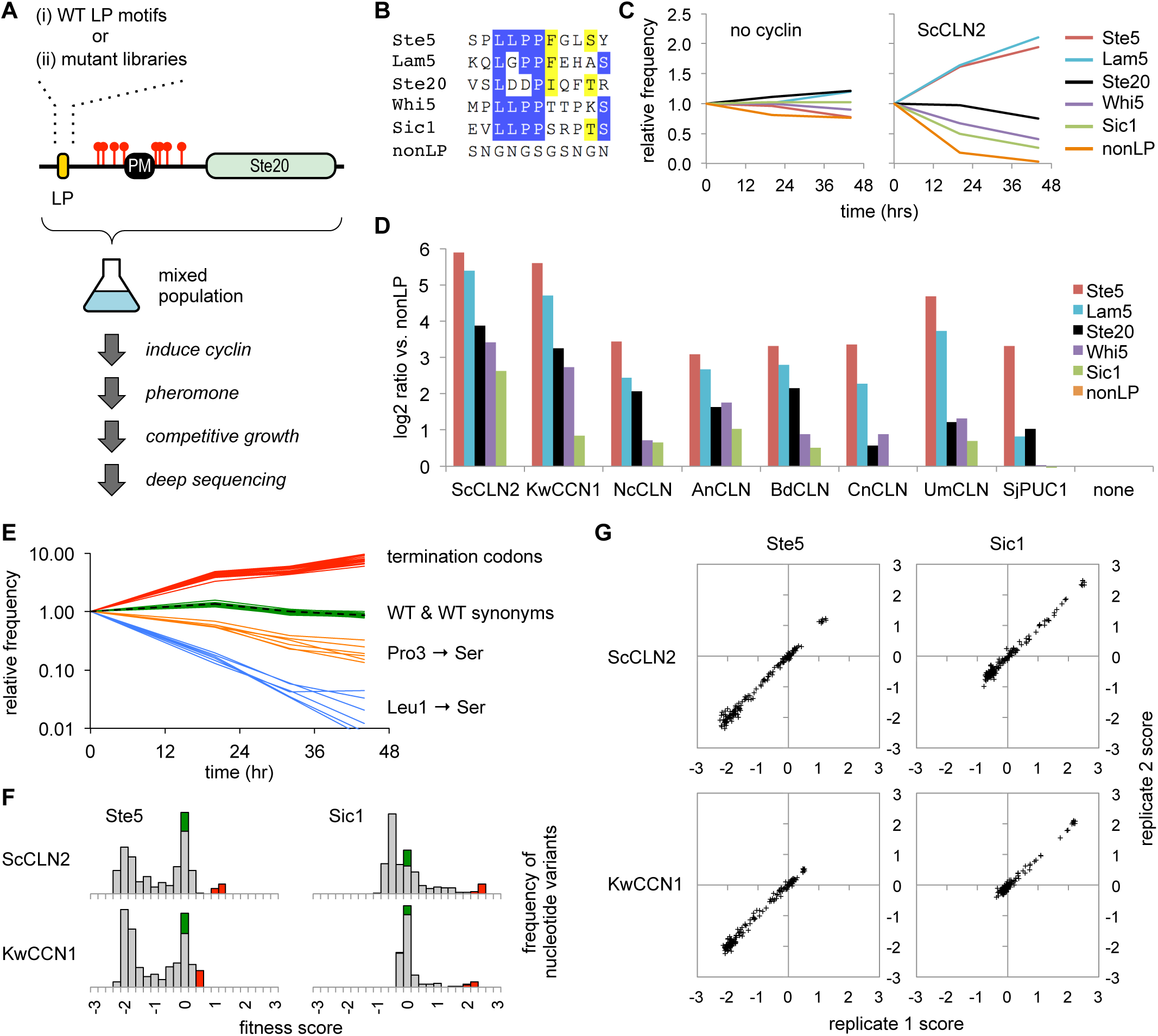
Competitive growth assay for the functional potency of cyclins and LP motifs. (A) Schematic diagram of the competitive growth assay, using Ste20^Ste5PM^ chimeras with a mixture of either wild-type (WT) LP motifs or libraries of randomized codon mutations. (B) Sequence comparison of five LP motifs and the control (nonLP) motif. (C) Examples of competitive growth results showing changes in frequency of 6 different LP motifs as a function of time after pheromone addition (0, 20, 44 hr), in cells with *P*_*GAL1*_*-ScCLN2* or no cyclin. Plots show averages of two independent experiments. (D) Experiments in panel C were performed in 8 different *P*_*GAL1*_*-cyclin* strains and the control strain (none). Bars show the ratio (log_2_) of the change in frequency for each motif compared to the nonLP control (at 44 hr), normalized to the results in the strain with no cyclin; values are the average of two independent experiments. (E) Example showing changes in frequencies of Ste5 LP motif variants (in *P*_*GAL1*_*-ScCLN2* cells). The dashed black line is the original WT sequence. Colored lines show all WT synonyms, all termination codons, and examples of missense synonyms with mild (Pro3 to Ser) or strong (Leu1 to Ser) defects. (F) Distribution of fitness scores for all nucleotide variants of two LP motifs (Ste5 and Sic1) in two *P*_*GAL1*_*-cyclin* strains. Green, WT synonyms; Red, terminators; grey, missense mutations. See Figure S1A for additional examples. (G) Correlation of fitness scores for all amino acid variants between two independent replicate experiments. See Figure S1B for additional examples.

Without a *P*_*GAL1*_*-cyclin* construct, none of the LP motifs had a notable selective advantage, but in strains expressing *ScCLN2*, they showed clear differences in enrichment and depletion (Figure 3C). The most potent motifs were those from Ste5 and Lam5, which became enriched over the control motif by roughly 7-fold after 20 hours and 50-fold after 44 hours. The next most potent was the motif from Ste20, followed by those from Whi5 and Sic1. A similar rank order was observed with G1 cyclins from seven other fungi (Figure 3D), though in some cases the weakest motifs were barely distinguishable from the nonLP control sequence; that is, cyclins that were more distant from *ScCLN2* were less able to utilize the weak motifs. These results reveal a range of functional potencies among different LP motifs that can lead to quantifiable growth differences, and indicate that the hierarchy of preference is generally conserved among divergent G1 cyclins. Strikingly, the strongest and weakest motifs contain the same core LLPP sequence, and hence the flanking residues must make important contributions to motif potency.

### Comprehensive analysis of LP motif sequence requirements

We exploited the competitive growth assay to analyze the determinants of LP motif potency in a comprehensive manner. First, the wild-type LP motifs in the Ste20^Ste5PM^ chimeras were replaced with sequences in which individual codons were randomized, to create libraries that substitute individual amino acids with all possible replacements (Figure 3A-ii). This was done for all 8 codons in each of the five LP motifs (Ste5, Lam5, Ste20, Whi5, Sic1). Then, the libraries were transformed into different *P*_*GAL1*_*-cyclin* strains and analyzed by the competitive growth assay, so that mutants with reduced or improved function would become depleted or enriched, respectively. The changes in population frequency for all mutants were analyzed using Enrich2 software [22], which calculates fitness scores based on their rate of depletion (negative scores) or enrichment (positive scores) relative to the wild-type motif (defined as zero).

All five LP motif libraries were analyzed multiple times in strains expressing *ScCLN2* and *KwCCN1*, plus additional trials with four other cyclins. Several general properties indicated that the behaviors of mutant variants were logical and reproducible: (a) Wild-type synonyms mimicked the original wild-type sequence (Figure 3E), with scores clustered around zero (Figures 3F, S1A), confirming that fitness levels reflected amino acid sequence irrespective of nucleotide sequence. Similarly, changes in fitness due to missense mutations were consistent regardless of the specific codon (Figure 3E). (b) Termination codons were the most favorable and thus yielded the highest scores (Figure 3E-F, S1A), which makes sense because they eliminate the protein required for the growth arrest signal. (c) Individual variants showed consistent scores in independent experiments (Figures 3G, S1B), indicating high reproducibility. (d) The largest negative scores were observed with the strongest motifs (e.g., Ste5, Lam5; Figure S1A-B), because they have the greatest difference compared to a non-functional motif (and so have the “most to lose”). By comparison, negative scores were smaller for weak motifs (e.g., Sic1, Whi5; Figure S1A-B), especially when combined with cyclins that poorly recognize the wild-type sequence (e.g., *KwCCN1* combined with the Sic1 motif).

The key features of each motif were revealed by plotting the comprehensive results as heat maps and sequence logos (Figures 4A-B). The Whi5 motif was unusual, as will be discussed later. For the other four motifs, several notable features emerged. (i.) Consistent with the invariant LxxP signature in all known motifs, there were strong preferences for Leu and Pro at positions p1 and p4, although Ala was tolerated surprisingly well at p4. (ii.) There was greater tolerance for substitution in the intervening positions, p2 and p3, with a clear hierarchy among the p2 residues found in wild-type motifs (i.e., Leu > Gly > Asp). (iii.) Despite a preference for Leu at p2, chemically similar residues such as Ile and Val were more disruptive than many dissimilar residues, such as polar or charged groups. (iv.) At p5 and p7 there were broader preferences for non-polar side chains, with Phe generally the most favored at p5. Notably, the preference at p7 was relatively mild with the strong motifs (Ste5, Lam5) and stronger with the weaker motifs (Ste20, Sic1). (v.) Positions p6 and p8 showed minimal selectivity, although a mild bias for polar groups at p6 was evident with the Ste5 motif. (vi.) With the Sic1 motif, mutations at p1 to p4 caused reduced fitness with *ScCLN2* but not with other cyclins (Figure 4A, S2), reflecting the fact that the wild-type motif is already nearly non-functional in combination with these other cyclins. Conversely, the Sic1 motif could be markedly improved (and hence recognized by all cyclins) by introducing non-polar residues at p5 or p7, indicating that the weak potency of the wild-type motif is caused by unfavorable residues at these positions.

**FIGURE 4:**
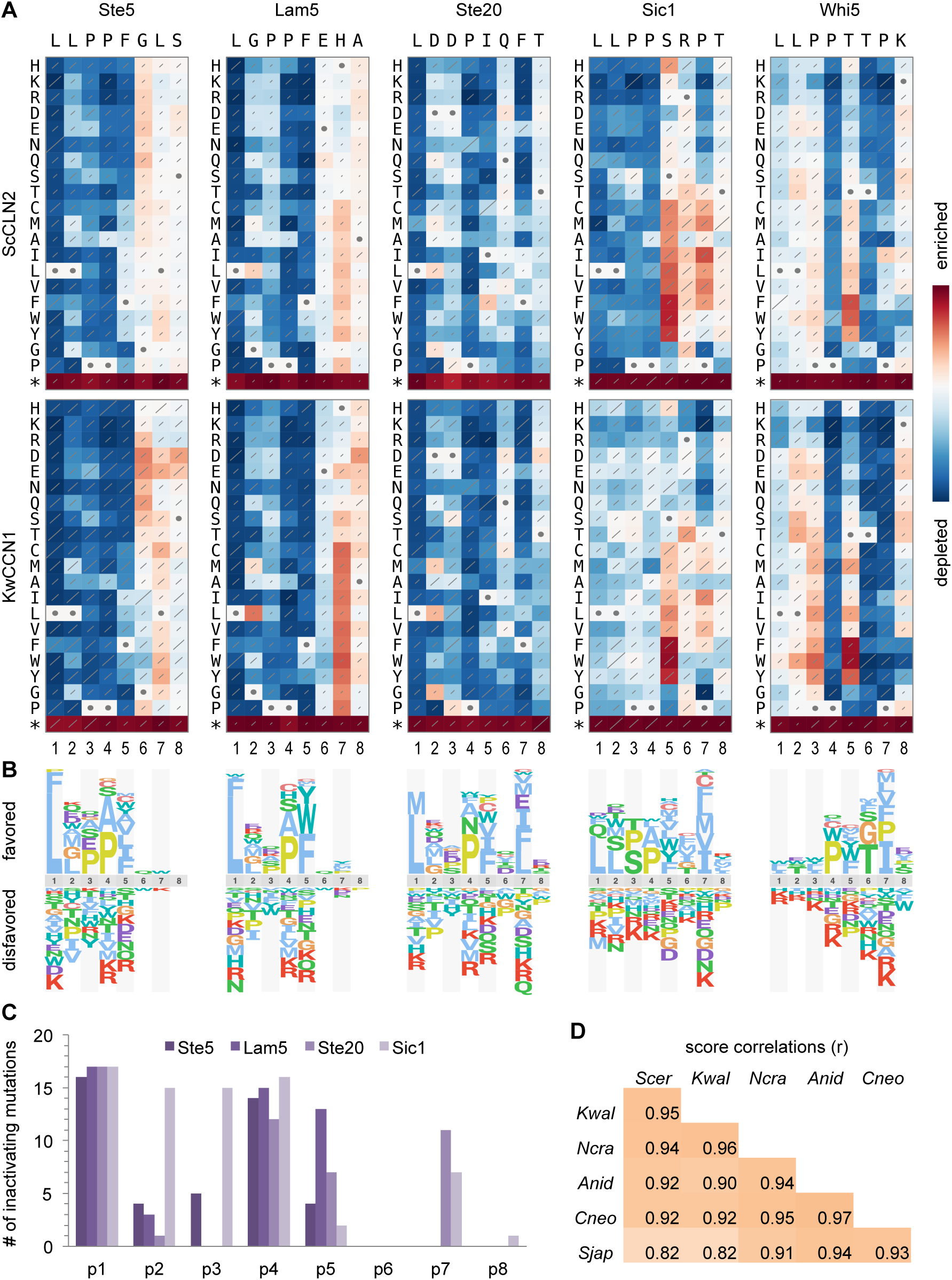
Comprehensive analysis of LP motif sequence preferences. (A) Heat maps show effects of all single-residue substitutions in 5 motifs. Red and blue indicate better and worse performance, respectively, relative to the wild-type motif. Each panel is individually scaled to its own maximum and minimum scores. Diagonal lines depict standard errors, scaled such that the highest value covers the entire diagonal. Circles denote the wild-type residue at each position. Data represent three independent experiments. See Figure S2 for analysis of the same motifs with 4 other cyclins. (B) Sequence logos showing relative preferences of *ScCLN2* for amino acids at each position of the motifs in panel A. (C) Plot of the number of inactivating mutations (defined as frequency scores within 15% of the minimum value; see Methods) at each position in 4 “typical” motifs (i.e., excluding Whi5). (D) Correlation coefficients (r) for all pair-wise comparisons of cyclins, calculated from the score matrices for all 5 motifs.

By calculating the number of inactivating mutations (Figures 4C, S3A), it was evident that p1 and p4 were generally the least tolerant of substitutions, followed by p5; the results at p2, p3, and p7 were more context-dependent, with p7 being less tolerant in the weaker motifs (Ste20, Sic1). A similar hierarchical pattern of selectivity of was also evident in measurements of functional inequality (Figure S3B), in which the different positions ranged from uniquely selective (p1, p4), to broadly selective (p2, p3, p5, p7) to nonselective (p6, p8). Importantly, the patterns of sequence preference were highly similar for all six cyclins tested (aside from their different capacities to recognize the Sic1 motif), including those from fungi beyond budding yeasts (*SjPUC1, AnCLN, NcCLN*, and *CnCLN*; Figure S2). Indeed, we found no notable examples of qualitatively distinct preferences. Instead, we found strong correlations for all pair-wise comparisons of cyclins (r = 0.82 to 0.97; Figure 4D), and in all cases the most favored residues fit the sequence LLPPΦxΦ (where Φ is hydrophobic). Overall, these results show that the LP motif sequence preferences are broadly conserved among fungal G1 cyclins. This finding is remarkable given that many of the known LP motifs, and even the proteins that harbor them, are not conserved beyond yeasts (see Discussion).

### Atypical behavior of the Whi5 sequence reveals overlapping motifs

Unlike other motifs, the Whi5 motif was not strongly impaired by mutations at p1 to p3 (Figure 4A). Moreover, it showed an extreme bias for a Thr residue at p6, a position that was not highly selective in other motifs. We considered two possible explanations for this atypical behavior. First, different flanking sequences might alter the reliance on positions within the 8-residue LP motif. Because the motifs were inserted as 11-residue segments with otherwise identical surrounding sequence, any such context effects would be limited to two preceding residues (positions p-2 and p-1) and one following residue (p9). Second, the required Thr residue at p6 is part of a TP dipeptide, a potential phosphorylation site. The resulting phospho-Thr moiety might improve cyclin binding, or it might be recognized by the Cks1 subunit of the cyclin-CDK complex [23, 24]; an overlapping Cks1 binding site could help explain why the Whi5 sequence is not fully reliant on the p1-p3 residues usually required for cyclin binding.

To address these possibilities, we repeated and extended the systematic analysis of four motifs (Ste5, Lam5, Sic1, Whi5) to include all 11 unique residues, and in parallel we analyzed two variant forms of the Whi5 sequence (Figure 5A). In one variant, S-Whi5, we altered residues flanking the 8-residue Whi5 motif to match those flanking the Ste5 motif, to test their influence on the different behaviors. In the other, called Whi5-P7L, we made a Pro to Leu substitution at p7, to simultaneously disrupt the potential TP phosphorylation site and improve the match to the p7 preference seen in other motifs. The results (Figure 5A) indicated minimal dependence on the flanking residues (p-2, p-1, and p9), as mutations at these positions had minor or variable effects (although a Pro residue was often disfavored at p-1). Furthermore, the altered flanking sequence in the S-Whi5 motif had no notable impact, as it retained the atypical behavior of the wild-type Whi5 motif. In contrast, the Whi5-P7L motif displayed radically altered behavior that essentially converted it from an atypical to a typical LP motif (Figure 5A): it showed increased dependence on positions p1 to p3 and a strong drop in preference for a Thr residue at p6. Thus, when the potential TP phosphorylation site was disrupted, the sequence reverted to typical LP motif preferences. These results support the notion that the Whi5 sequence contains an overlapping Cks1 site, which can promote CDK regulation of the substrate even when mutations (i.e., at p1 to p3) disrupt cyclin recognition of the LP motif. In this light it is noteworthy that the wild-type Whi5 motif showed a strong bias for a Pro residue at p4 (Figures 4A, 5A), which could imply a recognition requirement for either Cks1 or the kinase that phosphorylates the TP site. We also note that the relevant kinase could be either the CDK or the pheromone-activated MAPK (see Discussion).

**FIGURE 5:**
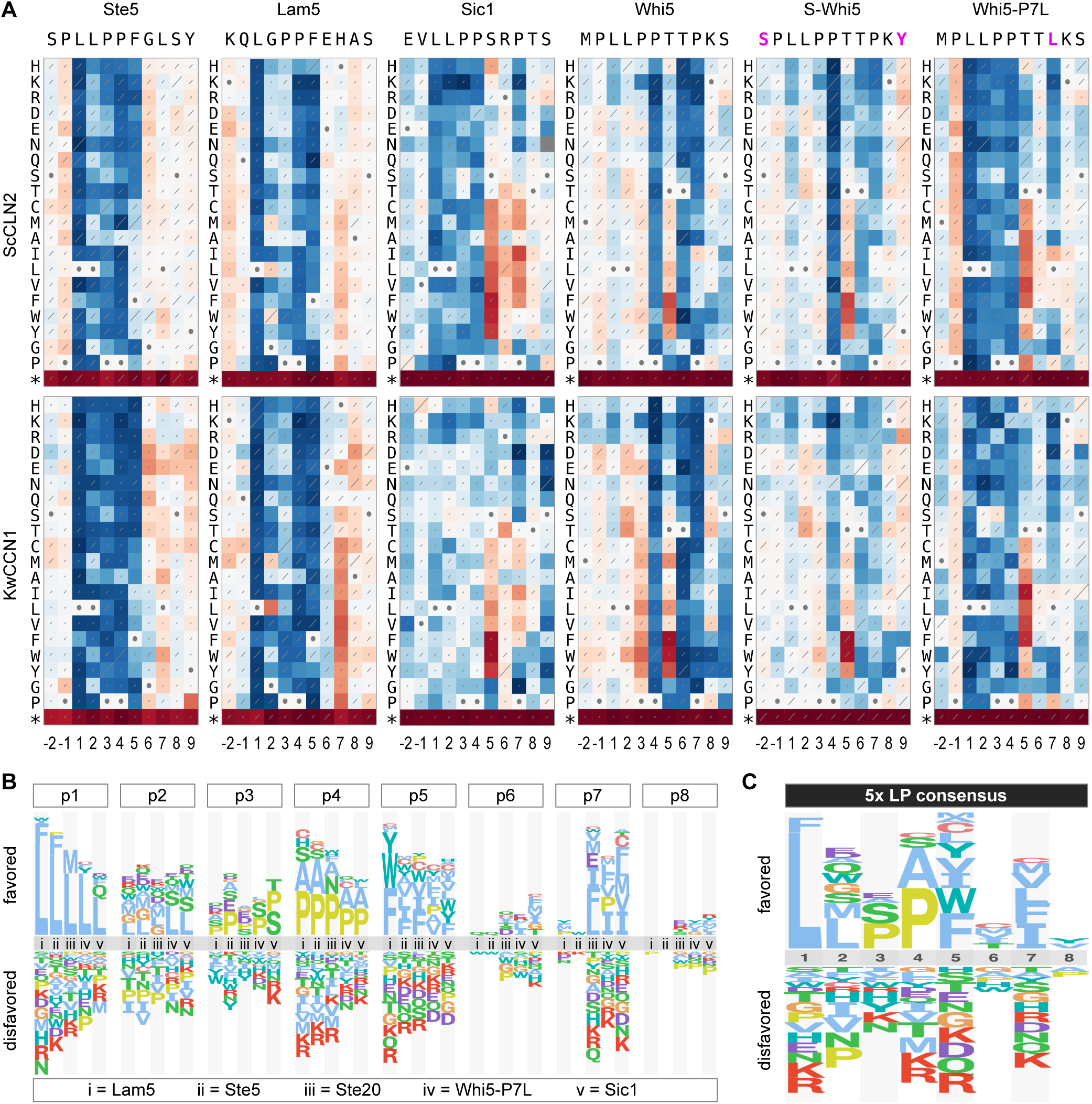
Analysis of atypical and consensus motif preferences. (A) Heat maps (as in Figure 4A) from mutational scanning of 11-residue sequences for 4 wild-type motifs plus two variants of the Whi5 motif (S-Whi5 and Whi5-P7L). Data represent two independent experiments. The Whi5-P7L results suggest that the atypical behavior of the Whi5 sequence is due to two overlapping motifs: an LLPP motif followed by a PxTP motif. (B) Sequence logos show the biases at individual positions, comparing five motifs with typical behaviors, including four wild-type motifs plus the Whi5-P7L variant. (C) Sequence logo showing the consensus preference averaged from the five motifs in panel B.

### A quantitative LP motif matrix predicts the potency of other candidate sequences

To distill the results from multiple distinct motifs into consensus preferences, we generated sequence logos that compared the biases at each position from the original four “typical” motifs (Ste5, Lam5, Ste20, Sic1) and the Whi5-P7L variant (Figure 5B). These plots highlighted the consistently strong selectivity at p1 (Leu), p4 (Pro or Ala), and p5 (non-polar), along with the context-dependent selectivity for non-polar residues at p7 that is most severe in weaker motifs (Ste20, Whi5-P7L, and Sic1). These quantitative preference scores constitute position-specific scoring matrices (PSSMs; see Methods) from each motif, which were averaged to generate a “consensus” PSSM (Figures 5C, S4A). We then used this consensus PSSM to objectively assess which amino acid properties might best explain the preferences at individual positions, by testing the scores for correlations to a prior set of 28 physicochemical descriptors [25]. The strongest correlations were seen at p2, p5, and p7 (Figure S4B). Positions p5 and p7 correlated best with the descriptor of “hydrophobicity”, in accord with the noted preference for nonpolar residues at these two positions. Position p2 correlated best with a complex descriptor (“steric hindrance / P[helix]”) that combines a preference for two properties – low steric hindrance and high α-helical propensity. This fits with the observation that, among chemically similar residues, proximal branches in side chains were disfavored (e.g., Leu > Ile/Val, Gln > Asn, Ser > Thr). Interestingly, the poorest correlations were observed at the most selective positions, p1 and p4, presumably reflecting a stringent requirement for a specific side-chain feature that is poorly mimicked by substitutes.

The preference for Pro residues at both p3 and p4 raised the possibility that the LP motif might adopt a polyproline II (PPII) helical structure. Because PPII helices show two-fold pseudosymmetry, peptide ligands that adopt this conformation can often present the key side chains equally well in either N-to-C or C-to-N orientation [26]. Thus, we compared four LP motifs in both forward and reverse orientation (Figure S4C), and we found that in all cases the forward orientation was strongly preferred. This finding helps constrain searches for candidate LP motifs in other proteins to a single orientation.

To further probe the sequence constraints that govern LP motif function, we tested several additional candidate motifs. These were selected from proteins that co-precipitate with Cln2 (unpublished observations) or are phosphorylated by Cln2-CDK [11, 17, 23, 27]. Within each protein, we located all LxxP motifs in regions predicted to be disordered, and then selected examples with varying degrees of matches to the favored residues at other positions (see Methods). We chose 14 examples to test by the competitive growth assay, in parallel with the Ste5 and nonLP motifs, using the full set of cyclins (Figure 6A-B). These sequences showed a broad range of potency, from stronger than Ste5 to indistinguishable from the nonLP control (Figure 6A), and their rank order was similar though not identical among the different cyclins (Figure 6B). We then used the consensus PSSM to calculate a predicted score for each motif, and found these scores correlated well with the observed results (R^2^ = 0.73-0.85; Figure 6C). Note that because all motifs contained a core LxxP sequence, their variation in potency must be due to the other positions. Overall, these findings indicate that the consensus PSSM derived from the systematic analyses can effectively predict the functional potency of new candidate motifs.

**FIGURE 6:**
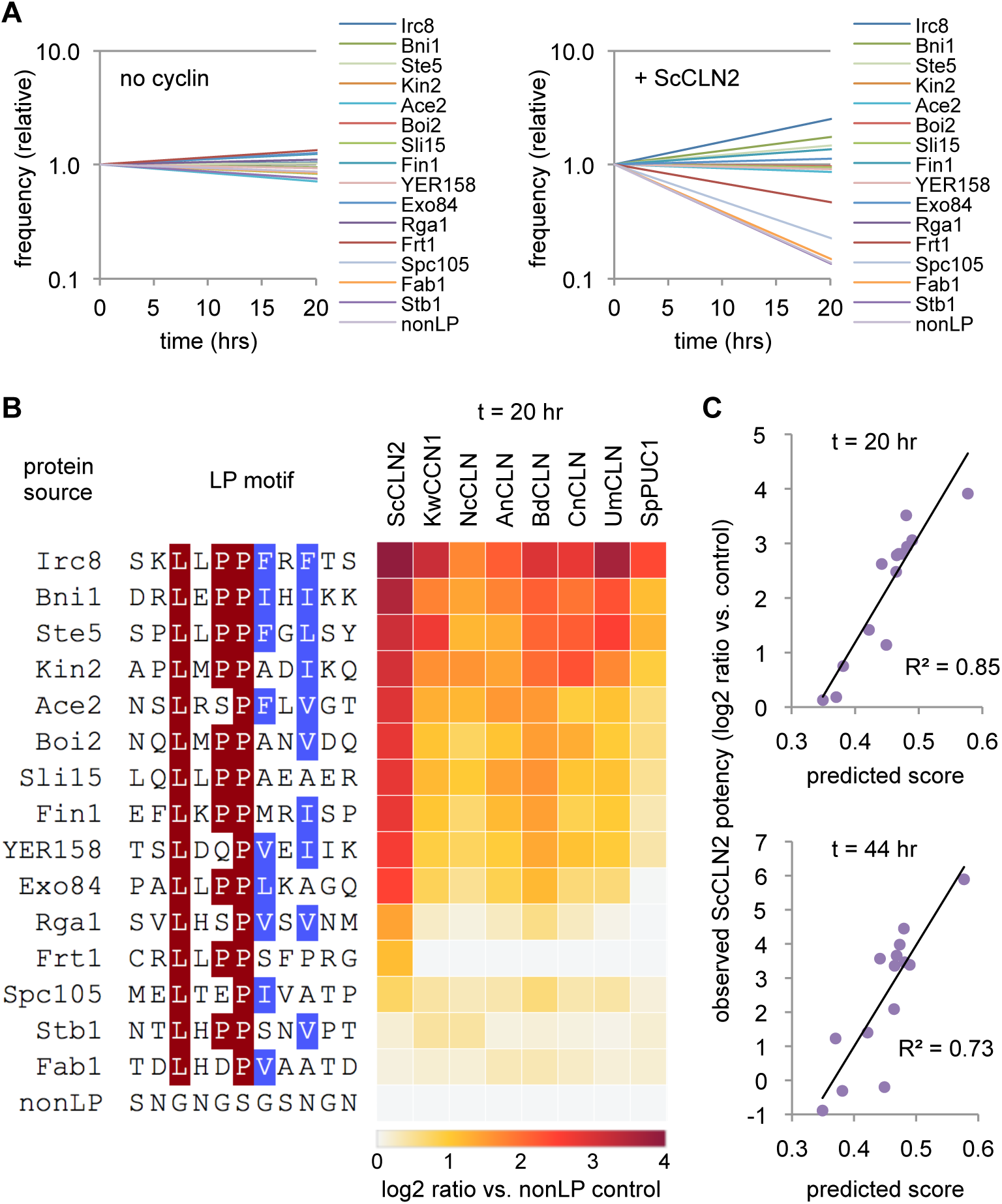
Observed and predicted potencies of additional candidate LP motifs. (A) Competitive growth results showing changes in frequency of 14 candidate LP motifs, plus the Ste5 and nonLP motifs, in cells with *P*_*GAL1*_*-ScCLN2* or no cyclin. Plots show averages of two independent experiments. (B) Heat map showing results of experiments in panel A performed with 8 different *P*_*GAL1*_*-cyclin* strains. Colors indicate the ratio (log_2_) of the change in frequency for each motif compared to the nonLP control (at 20 hr), normalized to the results in the no-cyclin strain; two independent experiments were averaged. At left are the source and sequence of each motif. (C) Comparison of observed vs. predicted potency for the 14 new candidate motifs (excluding the Ste5 and nonLP controls). Observed potency is the log_2_ ratio value (as in panel B) for the ScCLN2 strain, at both 20 and 44 hr. Predicted scores were derived from the consensus PSSM (see Figures 5C and S3A) by calculating the sum of preference score values at p1-p8 corresponding to the amino acid sequence for each LP motif.

### Disfavored residues in the Sic1 motif tune the timing of its CDK-mediated degradation

The LP motif from Sic1 showed weak potency with all cyclins. We reasoned that this weakness is due to the fact that, despite having an ideal LLPP sequence from p1 to p4, it lacks favored residues at positions p5 and p7. Indeed, replacement of either position with a hydrophobic residue caused an increase in motif potency (see Figures 4A, 5A). Curiously, we noticed that the absence of favored residues at p5 and p7 is generally conserved among Sic1 orthologs (Figure 7), raising the possibility that a weak LP motif in Sic1 is beneficial. In the native protein, this motif initiates a cascade of CDK phosphorylation reactions that ultimately trigger ubiquitin-mediated degradation of Sic1 and entry into S phase [28, 29]. Thus, we hypothesized that the submaximal potency of this LP motif helps set the timing of Sic1 degradation.

**FIGURE 7:**
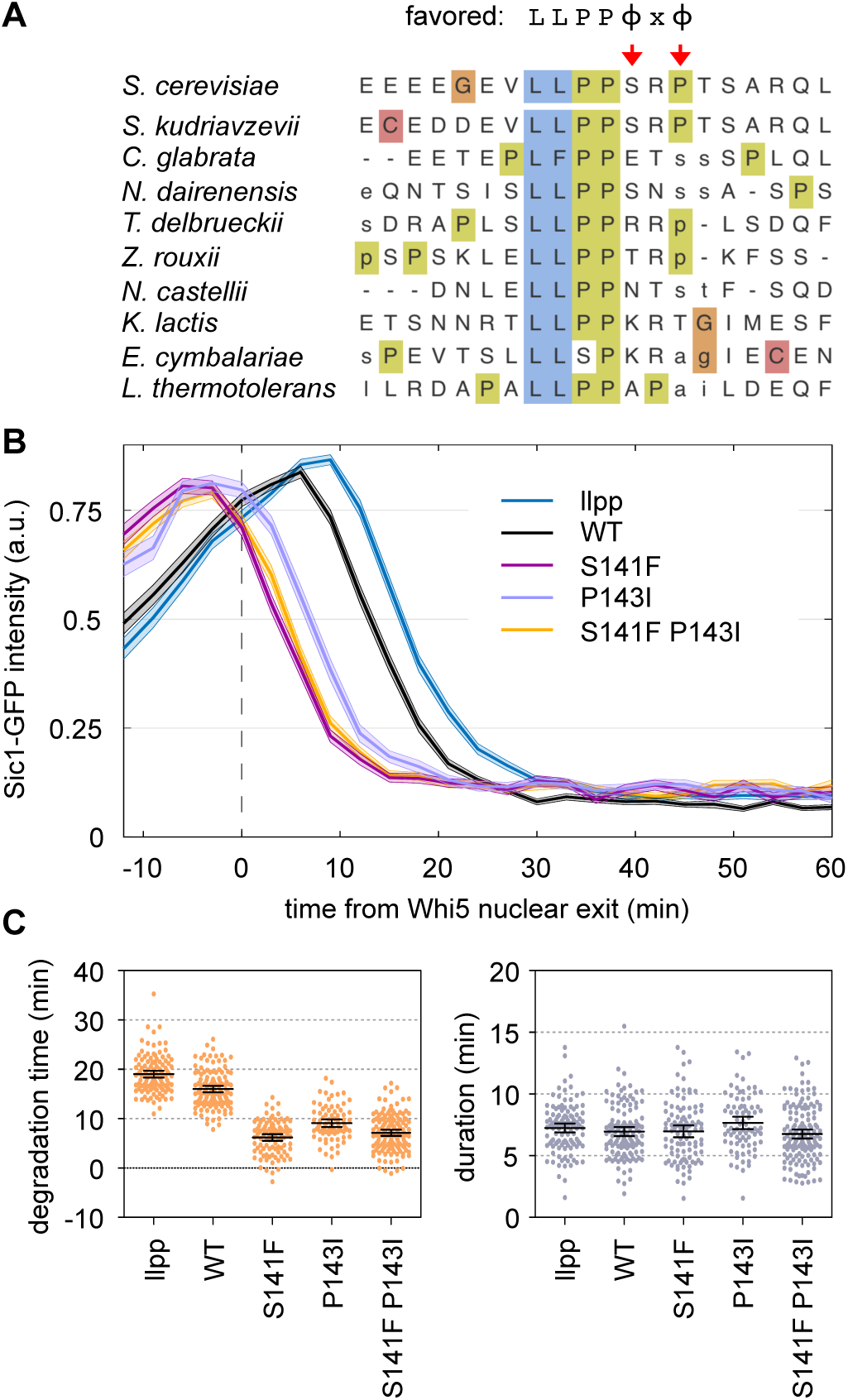
Unfavored residues in the Sic1 motif delay in its CDK-mediated degradation. (A) Alignment of the LP motif and proximal sequences from Sic1 orthologs in *S. cerevisiae* and nine other budding yeasts. At top is the consensus preferred residue pattern. Red arrows emphasize the conserved absence of favored residues at p5 and p7. (B) Plots of fluorescence intensity for variant forms of the Sic1-GFP reporter, relative to the time when nuclear exit of Whi5-mCherry is 50% complete in the same cells. Dark lines show the mean, and shaded bands show ± SEM. The llpp mutant substitutes VLLPP with AAAAA. Note that degradation timing was advanced by substituting either p5 or p7 with favored residues (S141F or P143I). (C) Degradation time and duration (see Methods) for the indicated Sic1-GFP reporters. Filled circles show results from individual cells, and black lines show the mean and 95% confidence intervals.

To test this view, we used a fluorescent reporter construct that allows Sic1 degradation to be monitored by live cell microscopy [13] (Figure 7B). Here, GFP is fused to a Sic1 N-terminal fragment (residues 1-215) that includes multiple CDK sites and the LP motif (other docking sites for later S-phase cyclins were removed). Within this construct, we changed the p5 and p7 positions of the LP motif to Phe and Ile, respectively, which yielded the strongest enhancements in the competitive growth assays. In native Sic1, these mutations correspond to S141F and P143I. Remarkably, each single mutation substantially accelerated the timing of Sic1-GFP degradation (Figure 7B-C), to a degree that was even greater than the difference between the wild-type motif and a defective motif (llpp). The effect was slightly stronger for S141F than for P143I (Figure 7B-C), which agrees with their relative strengths in the competitive growth assay, and the double mutation caused no further effect. The advanced timing caused the majority of Sic1-GFP to be degraded within 10 minutes after the cell cycle commitment point, or “Start” (Figure 7B); this is comparable to the earliest timing observed previously by adding an optimized di-phosphodegron [13]. Notably, this timing precedes the degradation of full-length, wild-type Sic1 and the consequent release of Clb-CDK activity [13, 17, 28, 29], and hence it is presumably driven solely by Cln1/2-dependent phosphorylation. We conclude that the disfavored residues in the Sic1 LP motif provide a beneficial tuning effect that delays the initiation of Sic1 degradation by Cln1/2-CDK until a time closer to the appearance of Clb-CDK activity, thus permitting the rapid two-stage relay of phosphorylation documented for Sic1 [28, 29].

## DISCUSSION

This study provides new insights into the mechanisms of substrate recognition by cyclin-CDK complexes. Although the use of LP motifs as docking sequences was originally discovered for the Cln1/2 cyclins [10, 28], we show here that this function is not restricted to budding yeasts and instead is an ancient property of fungal G1 cyclins that remains functionally intact in a broad range of extant species. By combining systematic mutagenesis and in vivo competition assays, we also uncovered the key sequence features that define LP motif function and potency. The maximally preferred features define a 7-residue consensus motif, LLPPΦxΦ, yet we also found varying degrees of plasticity at several key positions that allow motifs to vary in potency. Moreover, we observed that these variations in potency can provide a mechanism for tuning the sensitivity to CDK activity in order to dictate the cell cycle timing of substrate phosphorylation.

Our systematic analysis of motif sequence preference is analogous to approaches using peptide spot arrays, which have proven useful for analyzing binding or enzymatic activity of purified proteins in vitro [30, 31]. A beneficial aspect of our approach is that it operates under conditions present in living cells, including the presence of many competing interactions, and hence it intrinsically highlights variations in binding strength that are functionally consequential in vivo. Our analyses were also comprehensive, involving 34 combinations of cyclin and motif (i.e., 5 wild-type motifs combined with 6 cyclins, plus 2 variant Whi5 motifs combined with 2 cyclins). This expansive approach helped define the key requirements common to all motifs and cyclins, while also uncovering some features that showed context dependence; in particular, the preference for a hydrophobic residue at p7 was partly masked in strong motifs but was clearly pronounced in motifs that were weakened by disfavored residues elsewhere (i.e., p2, p3, or p5).

We found that variations in LP motif potency can help tune the magnitude and timing of regulation by Cln1/2-CDK. This notion was clearly illustrated for Sic1, where strengthening its LP motif advanced the timing of its degradation. The normal role of Sic1 is to prevent entry into S phase by blocking the appearance of Clb-CDK activity until the S-phase cyclin Clb5 has accumulated and the earlier Cln1/2-CDK activity has peaked [32, 33]; at this stage, Sic1 phosphorylation and degradation is initiated by Cln1/2-CDK and then is accelerated by the resulting release of Clb5-CDK activity [28, 29, 34, 35]. Our findings suggest that the weakness of the Sic1 LP motif helps delay the ability of Cln1/2 to initiate these events until Clb5-CDK complexes have accumulated and are poised for release from Sic1 inhibition. More generally, these findings emphasize that maximal affinity between SLiMs and their partners is not necessarily optimal. This view is reinforced by prior examples in which modulation of SLiM affinity could tune the balance of competing kinase and phosphatase activities on a substrate [4, 6]. Relatedly, although CDK enzymes prefer to phosphorylate SP/TP sites that are followed by basic residues, this feature is absent from the majority of CDK-regulated sites in yeast cells [36]. The fact that submaximal sequences can be functional, and even beneficial, poses a challenge to bioinformatic efforts to identify viable motifs in the proteome, and it cautions against demanding matches to a maximal consensus. This predicament emphasizes the utility of gaining comprehensive information of motif preferences, as in this study, that define not only favored residues but also prohibited substitutions that can provide filtering criteria to rule out nonfunctional sequences.

Surprisingly, the Whi5 sequence behaved as if it harbors two distinct overlapping motifs: an LP motif for cyclin binding and a potential phospho-Thr site for binding Cks1. Both types of binding sites can perform related roles in promoting substrate engagement and processive phosphorylation by CDK complexes [23]. As our experiments involved both pheromone treatment and cyclin expression, the kinase that phosphorylates the TP site could conceivably be either the CDK or the pheromone-activated MAPK. Indeed, we recently found that these two kinases can collaborate to regulate a shared target, Ste5 [37]. It should be noted that the dual-motif behavior of the Whi5 sequence observed in the context of the Ste20^Ste5PM^ chimera might not operate in the context of the native Whi5 protein. Yet it is also noteworthy that Whi5 orthologs show strong conservation of a 10-residue sequence (LLPPTTPKSR) that spans both the LLPP and TP sequences, plus C-terminal basic residues preferred by CDKs [38, 39].

Our findings indicate that the capacity for LP docking and the general motif preferences are conserved broadly in the fungal kingdom. Apart from yeast systems, fungal G1 cyclins have not been studied extensively, although recent work found that the *Aspergillus* Cln (PucA) has an essential role in the G1/S transition plus distinct non-essential roles [40]. In yeasts, in addition to cell cycle entry, Cln1/2-related cyclins promote polarized morphogenesis, as is evident during bud emergence in *Saccharomyces*, yeast-to-hyphae transitions in pathogenic *Candida*, and filamentous growth in *Ashbya* [41-44]. Hence, their counterparts in filamentous fungi might play similar roles during the polarized tip growth of hyphae, and LP docking could facilitate this role based on findings that it helps yeast Cln1/2 and Ccn1 cyclins induce polarized bud growth [17]. Because fungal G1 cyclins are thought to have evolved from a B-type cyclin precursor [18], it is likely that LP docking arose in fungi after their divergence from animals. Consequently, this function could present a target for anti-fungal therapeutics.

Despite the broad conservation, LP motifs are not recognized by Cln3 cyclins, suggesting they lost this function after diverging from Cln1/2. The function of Cln3 is highly specialized toward promoting early G1/S transcription; this is thought to involve inhibition of Whi5 [45, 46], yet Whi5 phosphorylation depends predominantly on Cln1/2-CDK [17], and so it is unclear if Cln3-CDK is merely a weak Whi5 kinase or if its key substrate is a distinct protein. Functional specialization of fungal G1 cyclins may be especially extreme in fission yeasts such as *S. pombe*, where *puc1Δ* mutants are viable [47]. Curiously, we found that LP motifs are recognized by Puc1 from *S. japonicus*. It is conceivable that Puc1 orthologs in these other fission yeasts have retained more evident roles, and hence it would be of interest to assess *puc1Δ* phenotypes in *S. japonicus*, especially given their capacity to alternate between yeast and filamentous morphologies [48].

Finally, although the ability of G1 cyclins to recognize LP motifs is widely conserved in fungi, the individual motifs are far less conserved. Indeed, for several cases in which LP motifs have been identified (e.g., Ste5, Sic1, Whi5), the proteins themselves lack obvious orthologs (or are highly diverged) outside of yeasts. As discussed previously [49], although SLiM-binding domains are often conserved over large evolutionary distances, the motifs can be rapidly acquired and discarded by target proteins. Because SLiM-binding domains often recognize motifs in dozens of other partner proteins, they face greater constraints to genetic drift than do the individual motifs. Furthermore, the key function of a particular SLiM interaction might often be achieved equally well by targeting any one of several proteins in the same pathway or multi-protein complex, which would allow for evolutionary drift in which protein harbors the SLiM. An example of this principle is illustrated by our ability to synthetically switch the CDK regulatory target in the pheromone pathway from Ste5 to Ste20. Therefore, additional efforts to comprehensively measure how the function and potency of SLiMs in vivo are altered by point substitutions, such as those that arise during genetic drift, will provide valuable insights into how evolution constructs and shapes SLiM-based regulatory networks.

## METHODS

### Yeast strains, media and growth conditions

Standard procedures were used for growth and genetic manipulations of yeast [50, 51]. Unless indicated otherwise, cells were grown at 30°C in yeast extract/peptone medium with 2% glucose (YPD) or in synthetic complete medium (SC) with 2% glucose and/or raffinose. Strains and plasmids used in this study are listed in Tables 1 and 2, respectively.

**TABLE 1:**
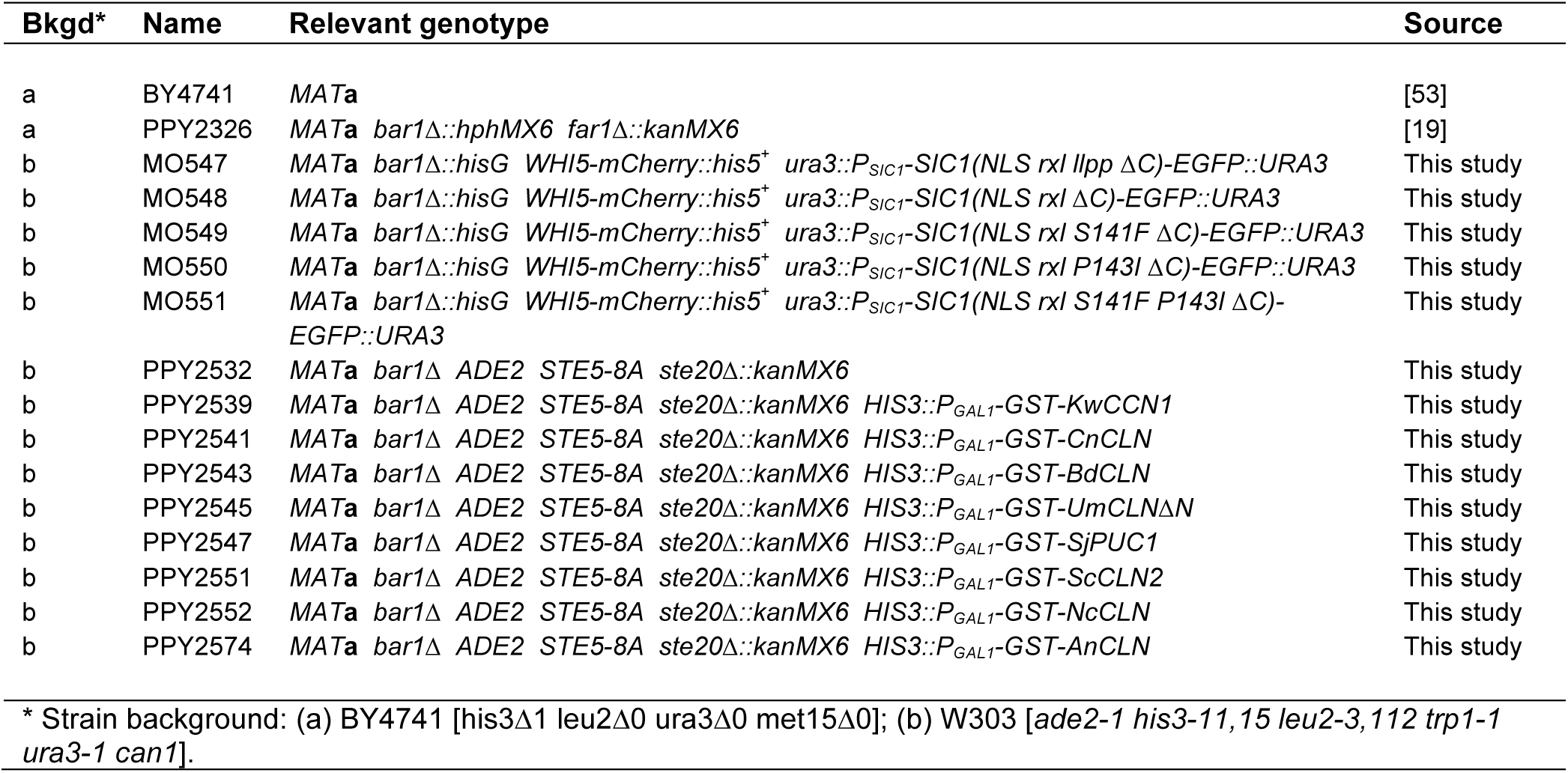
Yeast strains used in this study.

**TABLE 2:**
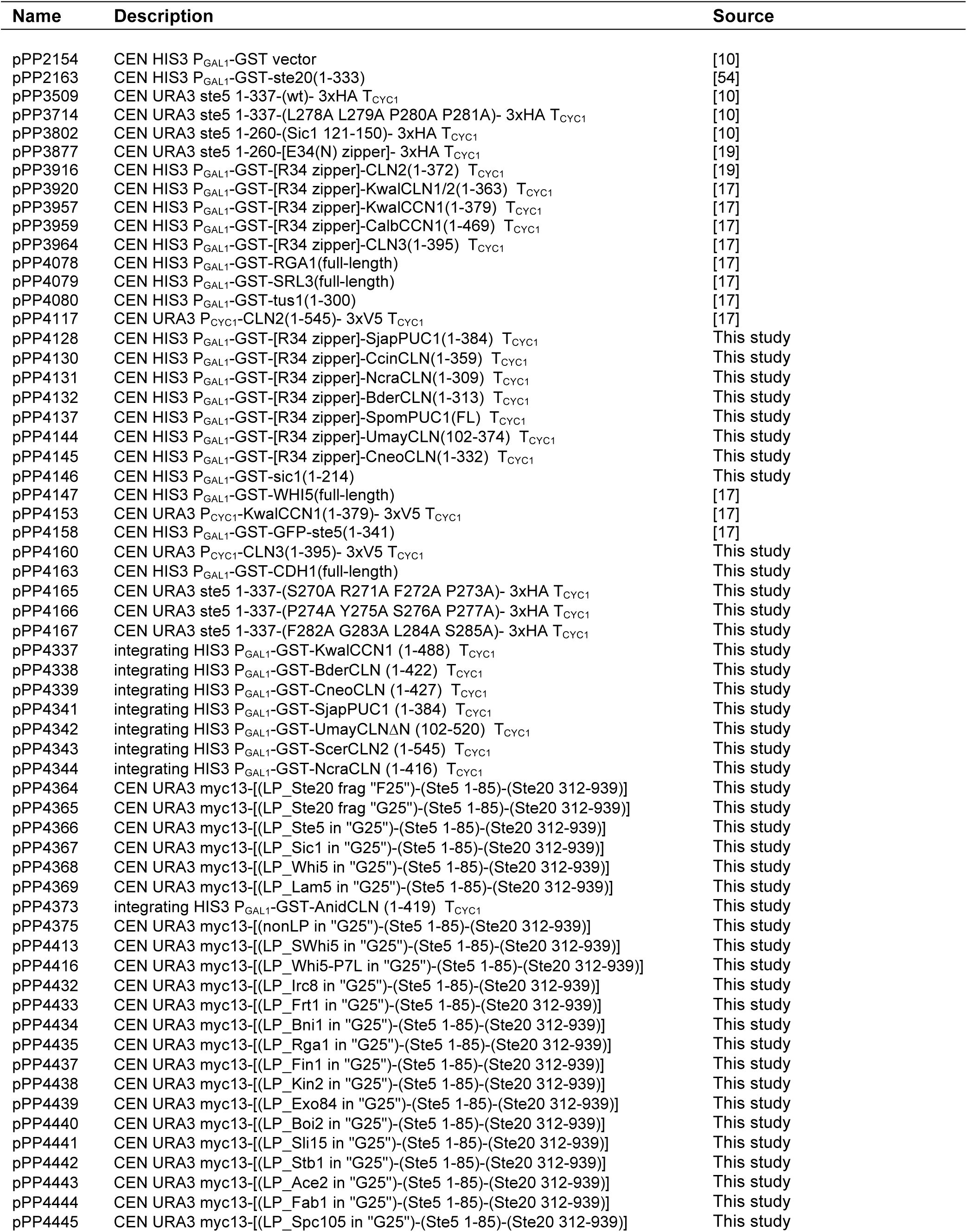
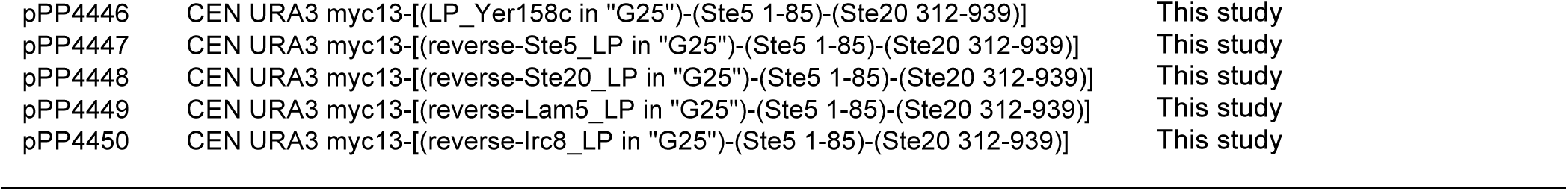
Plasmids used in this study.

Fungal G1 cyclin genes were amplified by PCR from genomic DNA samples and ligated into episomal or integrating *P*_*GAL1*_*-GST* vectors. For genes with introns (*AnCLN, NcCLN, BdCLN, CcCLN, CnCLN*), exon sequences were amplified separately and then joined together to create a pseudo-cDNA.

For mobility-shift assays of substrate phosphorylation, cells harbored plasmid-borne *P*_*GAL1*_*-GST-cyclin* expression constructs with C-terminal truncations (ΔC) that remove destabilizing sequences, which allowed maximal expression and accumulation of comparable protein levels for most cyclins. Full-length cyclins were used for pheromone-inhibition experiments and experiments involving longer-term expression, such as for in vivo selection of LP motif variants, because galactose-induced expression of the truncated forms inhibited growth. These full-length *P*_*GAL1*_*-GST-cyclin* constructs were integrated at the *HIS3* locus. Strains for microscopy expressed Whi5-mCherry from the native *WHI5* locus, and Sic1-GFP reporter constructs were inserted in single copy at the *URA3* locus, as described previously [13].

### Phosphorylation, protein-binding, and signaling assays

Mobility-shift assays to monitor cyclin-induced phosphorylation in vivo were performed as described previously [10, 17, 19]. Briefly, cells harboring *P*_*GAL1*_*-GST-cyclin* constructs and HA-tagged CDK substrates were induced with 2% galactose for 2.5 hr, to induce cyclin expression. Two mL of these cultures were harvested by centrifugation, and then the pellets were frozen in liquid nitrogen and stored at −80°C. As before (Ref_) all substrates were all based on the Ste5 N-terminus: (i) the “LP dock” substrate is Ste5 residues 1-337; (ii) the “no dock” substrate is Ste5 1-337 with its LLPP motif mutated to AAAA; (iii) the “leu zipper” substrate is Ste5 1-260 fused to a half leucine zipper sequence (E34[N]).

GST co-precipitation assays were performed as described previously [10, 17, 19]. Briefly, 10-mL cultures harboring *P*_*GAL1*_*-GST* fusion proteins and V5-tagged cyclins were treated with 2% galactose for 1.5 hr. Extracts were prepared by glass bead lysis in a nonionic detergent buffer, and then GST fusions and co-bound proteins were captured on glutathione-sepharose beads.

To assay the ability of fungal cyclins to inhibit pheromone signaling and MAPK activation, strains with integrated *P*_*GAL1*_*-GST-cyclin* constructs were grown to exponential phase in 2% raffinose media, and then cyclin expression was induced by adding 2% galactose for 75 minutes. Then, cells were treated with pheromone (50 nM) for 15 minutes. Two mL of these cultures were harvested by centrifugation, and then the pellets were frozen in liquid nitrogen and stored at −80°C.

### Whole cell lysates and Immunoblotting

Whole cell lysates were prepared from frozen cell pellets by lysis in trichloroacetic acid as described previously [19]. Protein concentrations were measured by a bicinchoninic acid assay (Thermo Scientific # 23225), and equal amounts (10 or 20 µg) were loaded per lane. Proteins were resolved by SDS–PAGE and transferred to PVDF membranes (Immobilon-P; Millipore) in a submerged tank. Membranes were blocked (1 hr, room temperature) in TTBS (0.2% Tween-20, 20 mM TRIS-HCl, 500 mM NaCl, pH 7.5) containing 5% non-fat milk, and then probed with antibodies in the equivalent solution. Primary antibodies were mouse anti-HA (1:1000, Covance #MMS101R), mouse anti-V5 (1:5000, Invitrogen #46-0705) mouse anti-phospho-p44/42 (1:1000, Cell Signaling Technology #9101), mouse anti-GST (1:1000, Santa Cruz Biotechnologies #sc-138), rabbit anti-myc (1:200, Santa Cruz Biotechnologies #sc-789), and rabbit anti-G6PDH (1:100000, Sigma #A9521). HRP-conjugated secondary antibodies were goat anti-rabbit (1:3000, Jackson ImmunoResearch #111-035-144), and goat anti-mouse (1:3000, BioRad #170-6516). Enhanced chemilluminescent detection used a BioRad Clarity substrate (#170-5060).

### Library construction and competitive growth assay

The construction of mutant libraries and the performance of bulk growth competition assays were based on previously described procedures [52]. LP motif variants were tested in the context of a chimeric Ste20^Ste5PM^ signaling protein [10], in which the membrane-binding domain from Ste5 and its flanking CDK phosphorylation sites replace the native membrane-binding domain in Ste20, and LP docking motifs are placed at the N-terminus. An initial parent construct (pPP4364) harbored the LP motif region from Ste20 on a 108-bp *Spe*I-*Bgl*II fragment that includes native Ste20 residues 80-109; in a later derivative (pPP4365), internal restriction sites were introduced within this region so that a smaller piece of the Ste20 LP motif sequence region (residues 86-96) could be replaced with 45-bp *Aat*II-*Nhe*I fragments that contain 11 unique codons between the *Aat*II and *Nhe*I sites (the products of which contain the descriptor “G25” in Table 2). To make libraries of point mutants, pPP4364 was digested with *Spe*I and *Bgl*II (for Ste20 libraries) or pPP4365 was digested with *Aat*II and *Nhe*I (for all other libraries). These digested vectors were ligated to annealed pairs of oligonucleotides (with single-stranded overhangs suitable for ligation with the vector) in which single codons were randomized (NNN) to generate all 64 possibilities. The ligation products were transformed into *E. coli* and plated on LB+Amp plates. Colonies (> 5000 per library) were harvested by adding LB+Amp liquid and gently agitating with a glass rod spreader, and then plasmid DNA was prepared from the suspension of pooled colonies. The isolated DNAs were checked by Sanger sequencing to verify that all four nucleotides were comparably represented at each position in the randomized codon.

For competitive growth assays, a solution was prepared that mixed together equimolar amounts of each individual codon library, plus a fraction (4-6%) of the wild-type plasmid. This pool was transformed into *P*_*GAL1*_*-cyclin* yeast strains. After the transformation procedure, an aliquot (10%) was reserved, and dilutions were spread on –URA plates; after 2 days incubation, colonies were counted to ensure that the number of transformants (usually > 10,000) exceeded the number of mutant variants in the library by at least 10-fold. The remainder of the transformation mixture was inoculated directly into 20 mL of –URA/glucose liquid medium and incubated at 30°C overnight (15-16 hr). Then, the cells were pelleted, washed five times with – URA/raffinose, and suspended in 50 mL of –URA/raffinose. This culture was incubated at 30°C for 48 hours, during which it was diluted back with fresh medium three times (at 22 hr, 31 hr, and 43 hr) to prevent saturation. The culture was then treated with 2% galactose for 75 minutes (in a volume of 70 mL) to induce cyclin expression. At this time (t = 0), an aliquot (40 mL) was harvested, and the remaining culture was treated with pheromone (500 nM) and returned to incubation. Cells were diluted with fresh medium (including galactose and pheromone) after the first 8 hours and subsequently after every 12 hours to maintain an OD_660_ below 1 (in a volume of 50 mL). Aliquots (20 mL) were harvested at 20, 32 and 44 hours. Harvested cells were collected by centrifugation (5 min., 5000 rpm, 4°C), washed with 25 mL sterile water, resuspended in 1 mL sterile water, and transferred to 1.5 mL microcentrifuge tubes. These suspensions were centrifuged, the supernatants were aspirated, and the pellets were frozen using liquid nitrogen and stored at −80°C.

### DNA preparation and deep sequencing

Isolation and sequencing of DNA from competitive growth assays followed previously described methods [52]. DNA was purified from yeast cells using the Zymo Research ZR Plasmid Miniprep Kit (#D4015). Frozen cell pellets were thawed and suspended in 200 µL of solution P1, and then were lysed using Zymolase (0.2 units/µL; Zymo Research #E1005) for 1.5 hours at 37°C, before proceeding with the remaining purification steps. The purified DNA was digested with *Not*I to improve the efficiency of the subsequent PCR. A test PCR (20 µL) was performed to confirm equal abundance of the template in all samples and to optimize the number of cycles for the final PCR (50 µL), which was 17 cycles for all results reported here. The forward primer included a P5 sequence (for binding the Illumina flow cell) followed by a Illumina sequencing primer binding site, a 6-nt bar code, and an upstream plasmid-annealing sequence; the reverse primer included a P7 sequence (for binding the Illumina flow cell) followed by a 6-nt i7-index sequence, an i7 sequencing primer binding site, and a downstream plasmid-annealing sequence.

Aliquots (5 µL) of the PCR products were run in 1.2% agarose gels to confirm the presence of the desired product, and the remainders were purified using Zymo Spin I columns (Zymo Research #C1003-250) and eluted in 10 mM TRIS-HCl, pH 8. The concentration of the eluted products was measured using nanodrop, and a mixture was prepared containing equal amounts of each DNA product. This final mixture was then re-purified using another Zymo Spin I column, and the DNA concentration was determined by qPCR using a KAPA Library Quantification Kit (Kapa Biosystems #KK4824). Samples were diluted to 20 nM, and 50 µL portions were submitted to the UMMS Deep Sequencing Core Facility for sequencing on an Illumina MiSeq instrument (single-end read, 100 bp, plus i7-Index sequencing when relevant).

Sequencing data were de-multiplexed by using bar code and index identifiers. Time-dependent changes in the frequency of individual sequence variants were analyzed using Enrich2 software [22]. Fitness scores calculated by Enrich2 describe the rate of change of mutant variants compared to the wild-type sequence, expressed as a natural log: i.e., (fitness score) = ln[(change in frequency of mutant variant) ÷ (change in frequency of wild-type)]. Overall, the relative fitness scores were similar regardless of whether they were calculated from the full time course (0, 20, 32, 44 hr) or just the initial stage (0-20 hr); all scores reported here were calculated from the 0-20 hr stage.

### Fitness score transformations and PSSM derivation

To create sequence logos of sequence preferences, we transformed the fitness score data generated by Enrich2, which is expressed relative to the WT motif sequence, into a preference metric that expresses the bias for each residue relative to the set of all possible residues. First, the raw fitness score for each amino acid variant was normalized to the lowest (= 0) and highest (= 1) scores in a given motif array: (normalized score) = (raw score – minimum) ÷ (mean terminator score – minimum). Then, the normalized scores were converted to a frequency metric by dividing each by the sum of all scores at the same position: (frequency) = (normalized score) ÷ (column sum). Finally, the frequency metric was converted to a preference metric by subtracting 0.05, so that a neutral preference is represented by zero, favored residues are positive, and disfavored residues are negative: (preference score) = (frequency – 0.05). To calculate the consensus PSSM, the individual PSSMs for 5 motifs (Ste5, Lam5, Ste20, Sic1, and Whi5-P7L) were averaged.

### Live cell microscopy assays of Sic1-GFP degradation

Time-lapse fluorescence microscopy of Whi5-mCherry localization and Sic1-GFP intensity was measured by methods that have been described in detail previously [13], using cells that were grown in SC/glucose medium, placed on agarose pads, and imaged at 30°C. Plots showing mean Sic1-GFP intensity were expressed relative to the time at which the nuclear Whi5-mCherry signal declined by 50%, and the Sic1-GFP degradation time in individual cells was defined as the point when its signal declined by 50%. Duration of Sic1-GFP signal was defined as the time from the start of its decline until the time it reached background levels.

## AUTHOR CONTRIBUTIONS

S. Bandyopadhyay performed experiments involving signaling inhibition and competitive growth assays. S. Bhaduri performed in vivo phosphorylation and co-immunoprecipitation assays. M.Ö. performed Sic1-GFP microscopy experiments, in the laboratory of M.L. N.E.D. analyzed fitness score data and generation of PSSMs. P.M.P. designed and oversaw the study, analyzed data, and wrote the manuscript.

## ACKNOWLEDGEMENTS

This work was supported by grants from the NIH (R01 GM057769) to P.M.P, by ERC Consolidator Grant 649124 and Estonian Science Agency grant PRG550 to M.L., and from CRUK (Senior Cancer Research Fellowship C68484/A28159) to N.E.D. We are indebted to Dan Bolon, Julia Flynn, and David Mavor for advice and guidance with deep mutational scanning methods. We also thank David Mavor and Alan Rubin for assistance with deep sequencing data processing and Enrich2 software, Matt Winters for technical help, Charlie Specht and Nick Rhind for fungal DNA samples, and Nick Rhind for helpful discussions.

---

**FIGURE S1:**
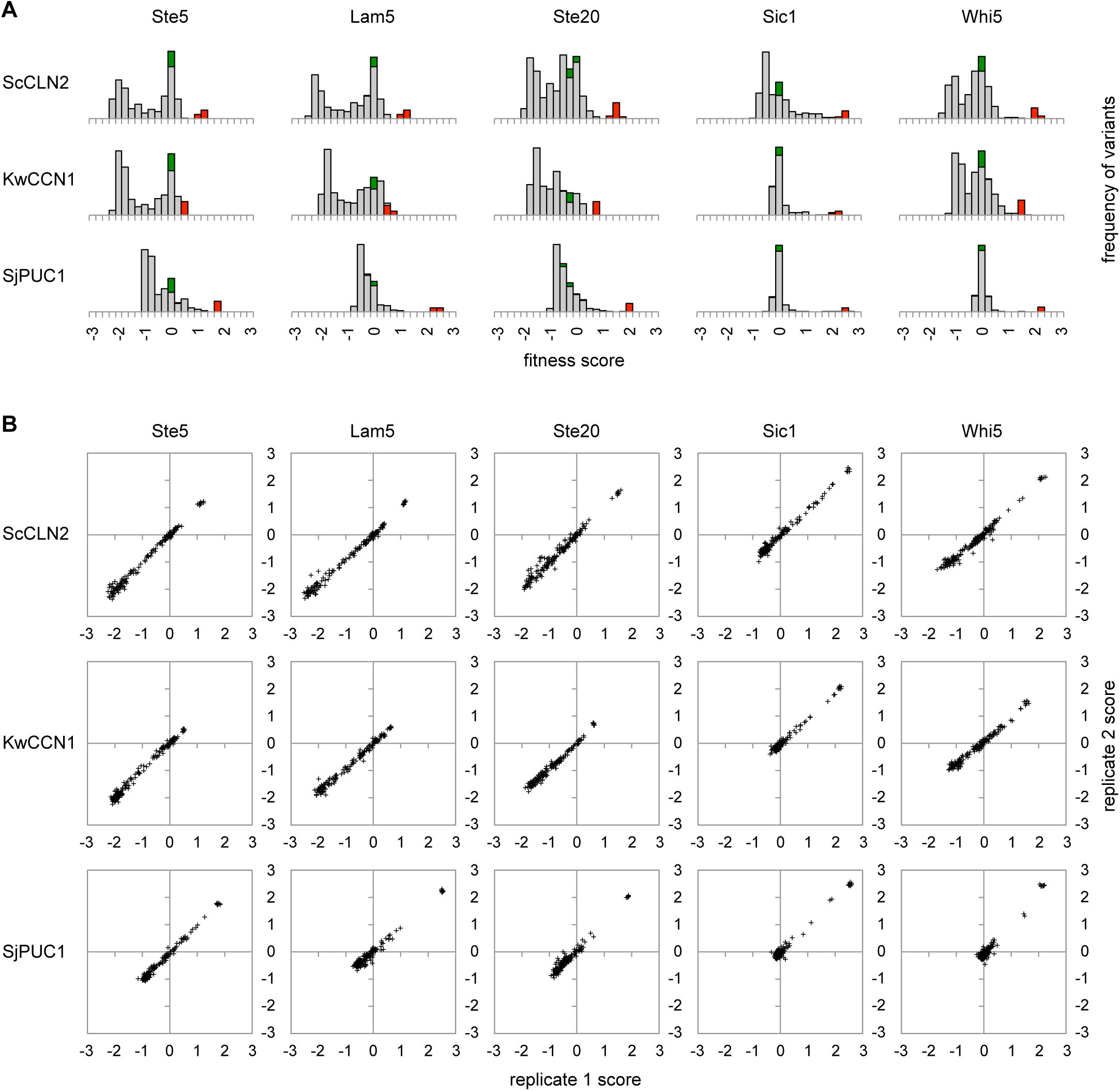
Score distributions and reproducibility for multiple cyclin-LP combinations. (A) Distribution of fitness scores for all nucleotide variants of all five LP motifs in three *P*_*GAL1*_*-cyclin* strains. Green, WT synonyms; Red, terminators; grey, missense mutations. Results are the averages for all experimental replicates (3 for ScCLN2 and KwCCN1, 2 for SjPUC1). (B) Correlation of fitness scores for all amino acid variants between two independent replicate experiments, shown for all five LP motifs and three cyclins.

**FIGURE S2:**
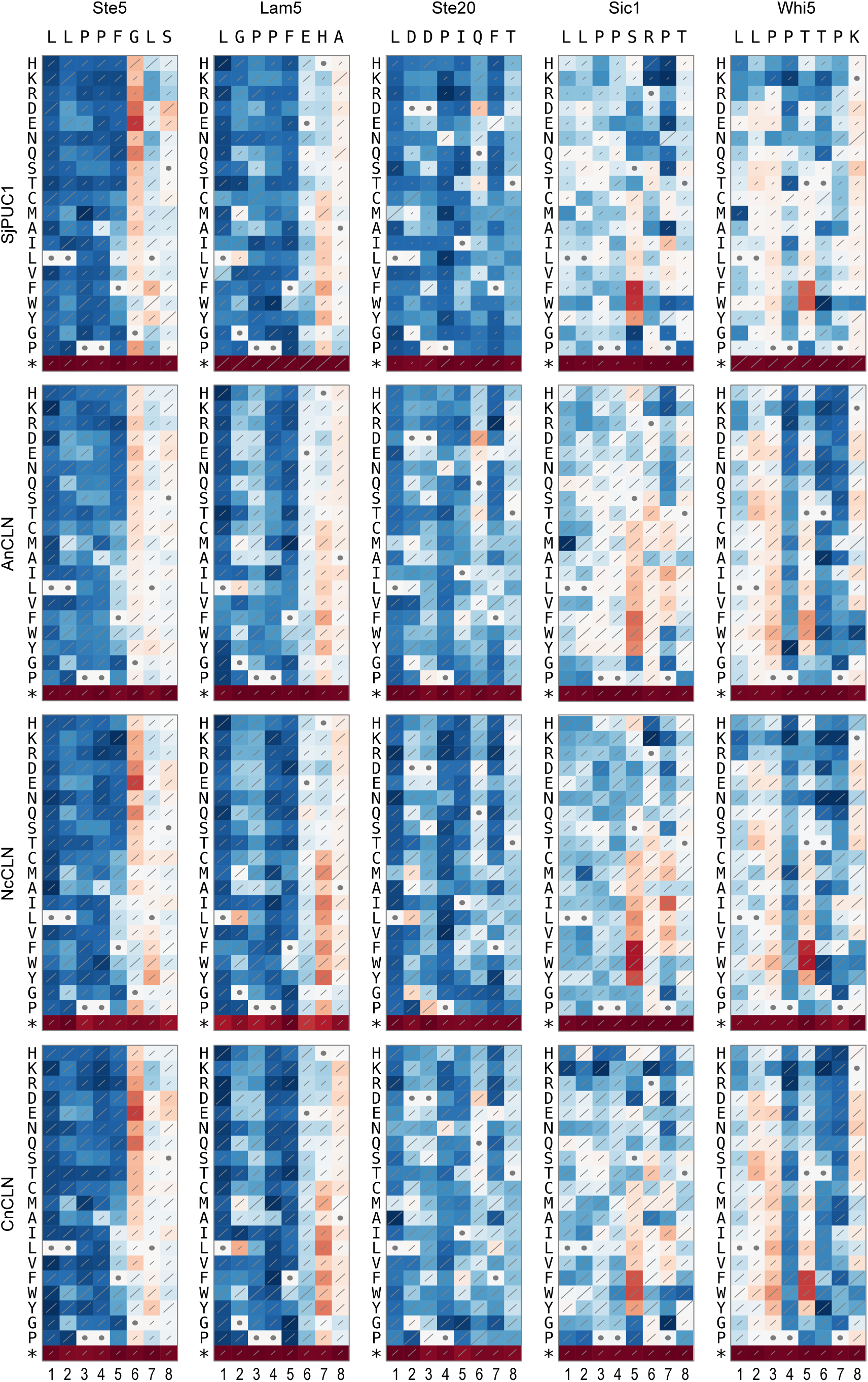
LP motif sequence preferences for cyclins outside of budding yeasts. Heat maps (as in Figure 4A) show all single-residue substitutions in 5 motifs affect recognition by cyclins from fission yeast and non-yeast fungi. Data represent two independent experiments for SjPUC1, or single experiments for AnCLN, NcCLN, and CnCLN (which are highly consistent with each other).

**FIGURE S3:**
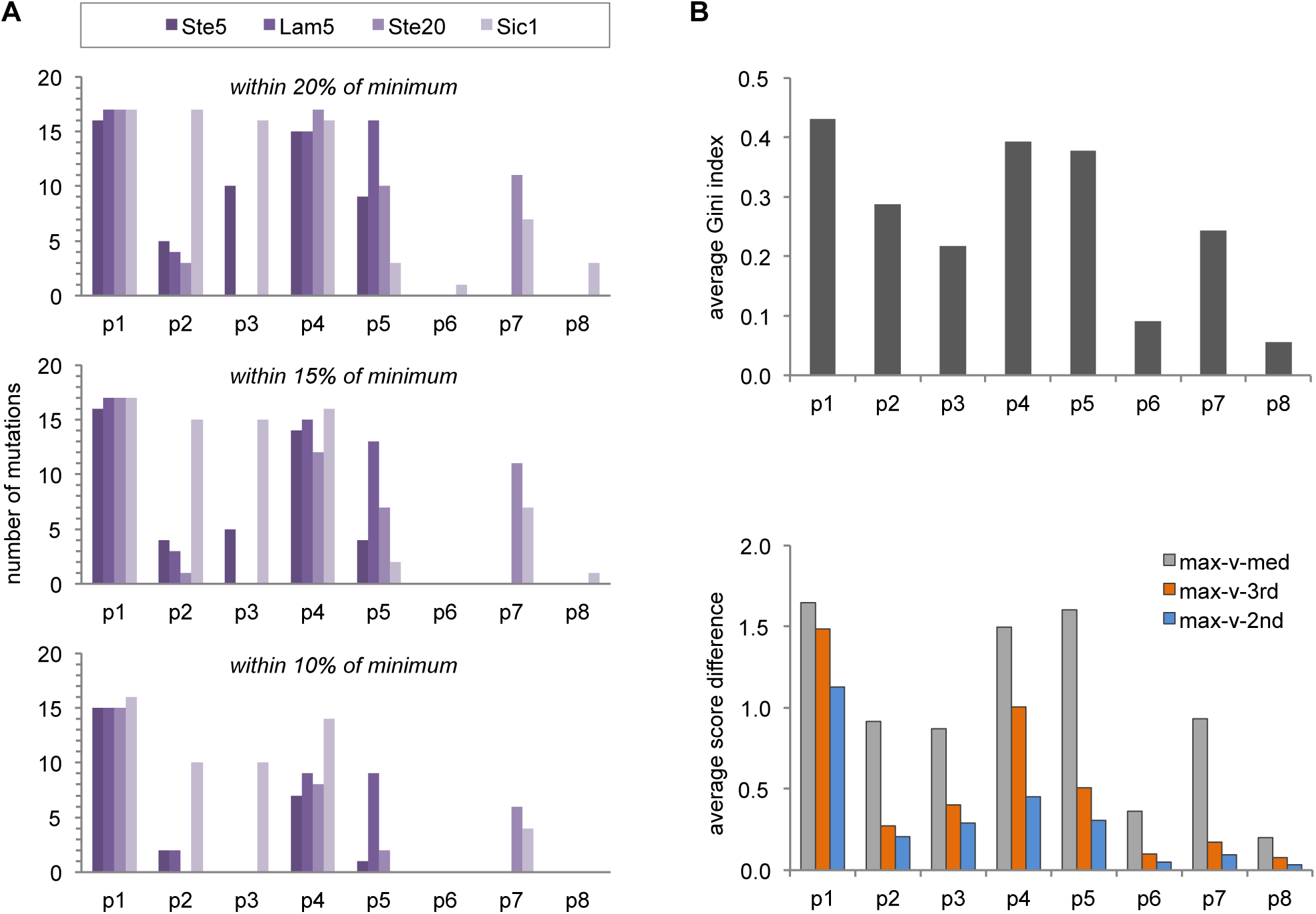
Degree of selectivity at different positions in the LP motif. (A) Comparison of the number of inactivating mutations using different threshold criteria. The plots show the number of frequency scores (see Methods) that were within 20%, 15%, or 10% of the minimum value, at each position in the 4 “typical” motifs (i.e., excluding Whi5). The results for the 15% criterion are shown in Figure 4C. (B) *Top*, plot of the Gini index (averaged for the 4 non-Whi5 motifs), which measures the degree of inequality in scores among the set of all possible residues. *Bottom*, plots of the difference between the maximum fitness score (max) and the median score (med), the 3rd highest score (3rd), and the 2nd highest score (2nd), averaged for the 4 non-Whi5 motifs. Positions that are highly selective for one or two unique residues (e.g., p1 and p4) show large differences between maximum and the 3rd and/or 2nd scores, whereas positions with broader preference for a group of residues (e.g., p5 and p7) show large differences between maximum and median but not between maximum and 3rd and/or 2nd scores.

**FIGURE S4:**
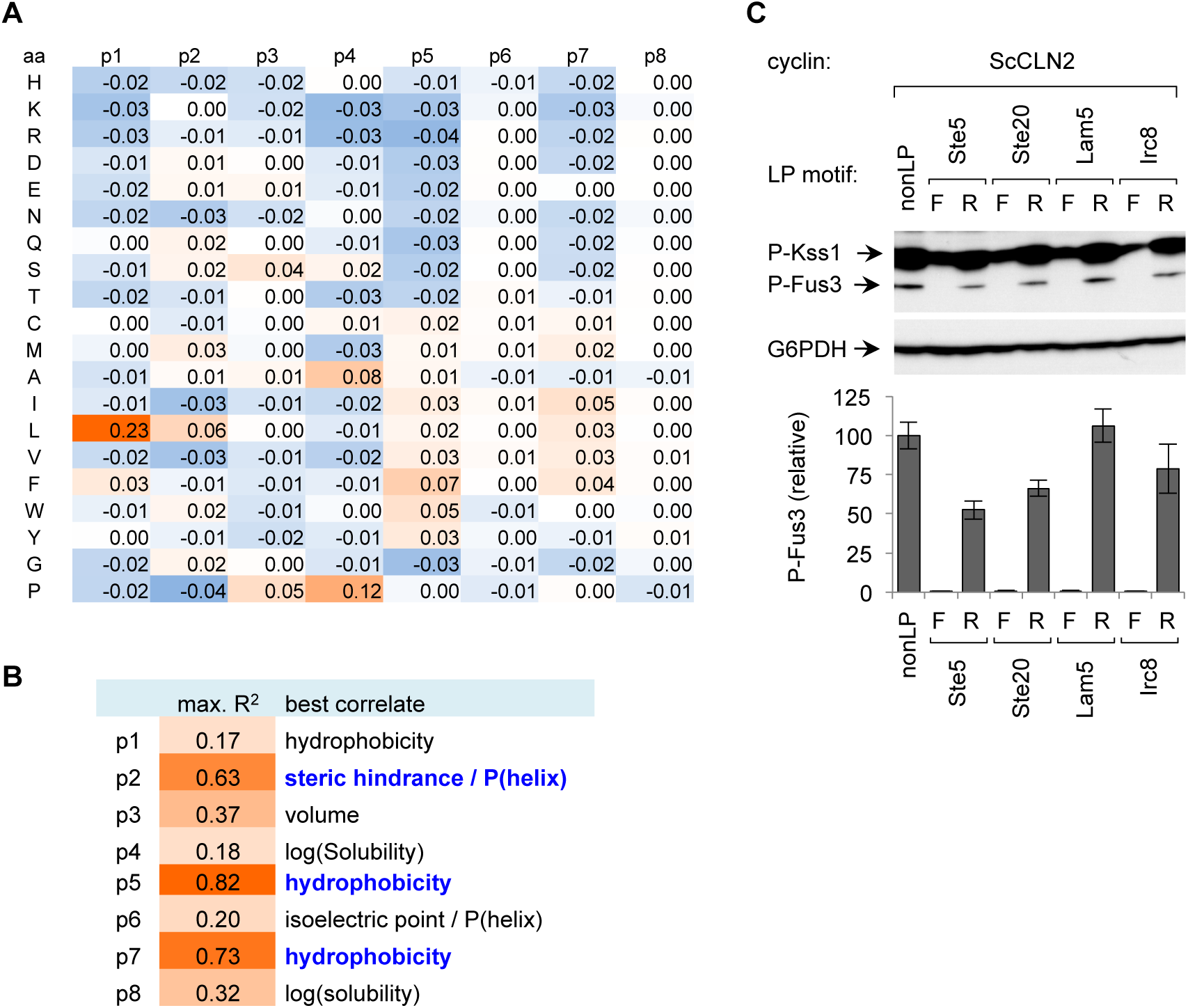
Consensus PSSM properties and orientation dependence of motifs. (A) Consensus PSSM, obtained by calculating the average of individual preference score PSSMs for 5 motifs (Ste5, Lam5, Ste20, Sic1, and Whi5-P7L; see Methods). (B) Comparison of amino acid properties with their preference patterns thoughout the LP motif. The values at each of the 8 positions in the consensus PSSM were tested for correlation to 26 physico-chemical descriptors [25]. The descriptor with the strongest correlation, and its R-squared value, are show for each position. Bold, blue text indicates the strongest correlations. (C) Orientation-dependence of LP motif sequences. Ste20^Ste5PM^ chimeras were constructed that contained LP motif sequences in either forward (F) or reverse (R) orientation. *P*_*GAL1*_*-ScCLN2* cells harboring these chimeras were treated with pheromone (50 nM, 15 min.) and tested for MAPK phosphorylation. Below are quantified results (mean ± range, n = 2).

